# Neutralization Assay with SARS-CoV-1 and SARS-CoV-2 Spike Pseudotyped Murine Leukemia Virions

**DOI:** 10.1101/2020.07.17.207563

**Authors:** Yue Zheng, Erin T. Larragoite, Juan Lama, Isabel Cisneros, Julio C. Delgado, Patricia Slev, Jenna Rychert, Emily A. Innis, Elizabeth S.C.P. Williams, Mayte Coiras, Matthew T. Rondina, Adam M. Spivak, Vicente Planelles

## Abstract

Antibody neutralization is an important prognostic factor in many viral diseases. To easily and rapidly measure titers of neutralizing antibodies in serum or plasma, we developed pseudovirion particles composed of the spike glycoprotein of SARS-CoV-2 incorporated onto murine leukemia virus capsids and a modified minimal MLV genome encoding firefly luciferase. These pseudovirions provide a practical means of assessing immune responses under laboratory conditions consistent with biocontainment level 2.

## Introduction

Coronaviruses are a group of enveloped RNA viruses with a positive-sense single-stranded RNA genome ranging from 26-32 kilobases, which can cause respiratory tract infections. In December 2019, a novel coronavirus known as severe acute respiratory syndrome coronavirus 2 (SARS-CoV-2) was identified in China and has caused a global ongoing pandemic of coronavirus disease (COVID-19). To date, SARS-CoV-2 has spread to 188 countries (https://coronavirus.jhu.edu/). More than 29 million cases and 900,000 deaths have been reported at the time of this writing.

Enveloped viruses are known to efficiently package their core elements with heterologous envelope glycoproteins, giving rise to the so called ‘pseudotypes’ or ‘pseudoviruses’. Many laboratories have successfully generated pseudotypes containing the core elements of HIV-1 [1] or MLV [2, 3] and the envelope glycoproteins of vesicular stomatitis virus [4], murine leukemia virus [5], Lassa fever virus, ebola virus, coronavirus spike glycoproteins, and others (reviewed in [6]).

In a pseudotype virus, viral attachment [7], entry, and importantly, antibody binding and neutralization sensitivity are dependent on the membrane glycoprotein provided [6]. Using a defective MLV vector genome encoding *firefly* luciferase, and a packaging vector encoding MLV gag/pol, we describe the production of pseudovirus particles containing the spike glycoprotein of SARS-CoV-2. As controls, we also produced similar particles containing SARS-CoV-1, VSV-G or HIV-1 LAI gp160.

## Materials and Methods

### Cells

HEK293FT cells, Vero E6 cells, SupT1 cells and Huh7 cells were purchased from ATCC. HEK293FT, Vero E6 and Huh7 cells were cultured in Dulbecco’s modified Eagle’s medium (DMEM) (Gibco, US) supplemented with 10% FBS (Gibco, US) and 2mM L-glutamine (Gibco, US) at 37°C with 5% CO2. 293ACE2 cells were cultured in DMEM with 10% FBS, 2mM L-glutamine and 200ug/ml hygromycin B (ThermoFisher, US).

### Plasmids

SV-Psi^-^-Env^-^-MLV [8], pHIV-1 LAI gp160 [9], pHCMV-VSV-G [4] and pSIVmac gp130 [10] were previously described. L-LUC-SN was constructed by inserting the *firefly* luciferase gene within the polylinker of pLXSN (Clonetech, cat# 631509). pSARS-CoV-1 was purchased from Sino Biologicals. pCAGGS expressing SARS-CoV-2 RBD was obtained from BEI Resources (cat#NR-52309). HEK293T-hACE2 cells were a gift from Adam Bailey and Emma Winkler and were constructed as follows. A DNA fragment containing a codon-optimized version of hACE2 (Genbank NM_021804) was inserted into pLV-EF1a-IRES-Hygro (Addgene Plasmid #85134) using Gibson assembly. 293T cells were then transduced with lentivirus made from this construct. The plasmid pcDNA3.1-SARS-2-S-C9 was a generous gift from Tom Gallagher and expresses a codon-optimized SARS-CoV-2 spike open reading frame with a deletion in the 19 carboxy-terminal deletion amino acids (an endoplasmic reticulum retention signal) and addition of the C9 peptide TETSQVAPA, recognized by antibody 1D4.

### Production of pseudotyped MLV

The plasmid SV-Psi^-^-Env^-^-MLV and L-LUC-SN were co-transfected with or without an envelope glycoprotein plasmid (pHCMV-VSV-G/pSARS-CoV-1/pSARS-CoV-2/pHIV-1 LAI gp160) into HEK293FT cells using Lipofectamine™ 3000 (ThermoFisher, US). Cell supernatants containing viruses were collected after 2 days of transfection. Viruses were filtered through a 0.45μm filter (VWR, US) and centrifuged at 4°C, 6500rpm for 18h over a 20% sucrose cushion. Viruses were resuspended in 500μl cell culture medium and stored at -80°C.

### Pseudovirus infection

HEK293FT, 293T-ACE2, and Huh7 cells were seeded in 96-well plates (ThermoFisher, US) the day before infection. SupT1 cells were added into a 96-well plate at the time of infection. 5⨯10^4^ cells were added to each well. Pseudotyped MLV viruses were added to the pre-cultured cells. Cells were cultured at 37°C with 5% CO_2_ for 2 days. All cells in each well were lysed and luciferase was measured using ONE-Glo™ Luciferase Assay reagent (Promega, US). RLUs are per well of a 96-well plate.

### Neutralization assay

293T-ACE2 cells were seeded in 96-well plates at 5⨯10^4^ cells per well the day prior to infection. Sera from COVID-19-positive patients, negative sera, positive control (RBD) and negative control (SIVgp130) were serially diluted in a volume of 100μL and pre-incubated with 50μL of pseudotyped viruses at 37°C for 1h. For these infections, virus stocks were used at a dilution resulting in 100-200 RLU in the absence of serum. Cells were then infected with the serum/pseudovirion mixtures. Luciferase was measured 48 hours post infection using ONE-Glo™ Luciferase Assay reagent. Neutralization titers NT_50_ and NT_80_ were calculated using Prism 8 (GraphPad, US).

## Results

To generate pseudovirion particles, three plasmids were co-transfected into HEK293FT cells. The first plasmid was the packaging construct, SV-Psi^-^-Env^-^-MLV; the second plasmid was L-LUC-SN, a minimal retroviral transfer vector encoding the *firefly* luciferase reporter gene; the third plasmid was an expression construct encoding one of the following membrane viral glycoproteins: SARS-CoV spike (hereafter referred to as SARS-CoV-1), SARS-CoV-2 spike, HIV-1 LAI gp160 and VSV-G. VSV-G pseudotyped virus is used as a positive control because of its high infectivity in most cell types. HIV-1 LAI gp160-pseudotyped virus is used as a negative control as it utilizes CD4 as a primary receptor, which is present in SupT1 cells but absent in HEK293T.

Pseudotyped MLV viruses were tested on HEK293FT, HEK293T-ACE2, Huh7 and SupT1 cells. HEK293FT cells were used as a control cell line, which is known to lack of susceptibility of coronavirus and HIV due to the absence of both ACE2 and CD4. As expected, VSV-G pseudotyped viruses infected all cell types and showed the highest infectivity (Figure 1). HIV-1 LAI gp160-pseudotyped viruses only infected SupT1 cells. Both SARS-CoV-1 spike pseudotyped virus and SARS-CoV-2 spike pseudotyped viruses infected 293T-ACE2 and Huh7 cells.

**Figure 1.**
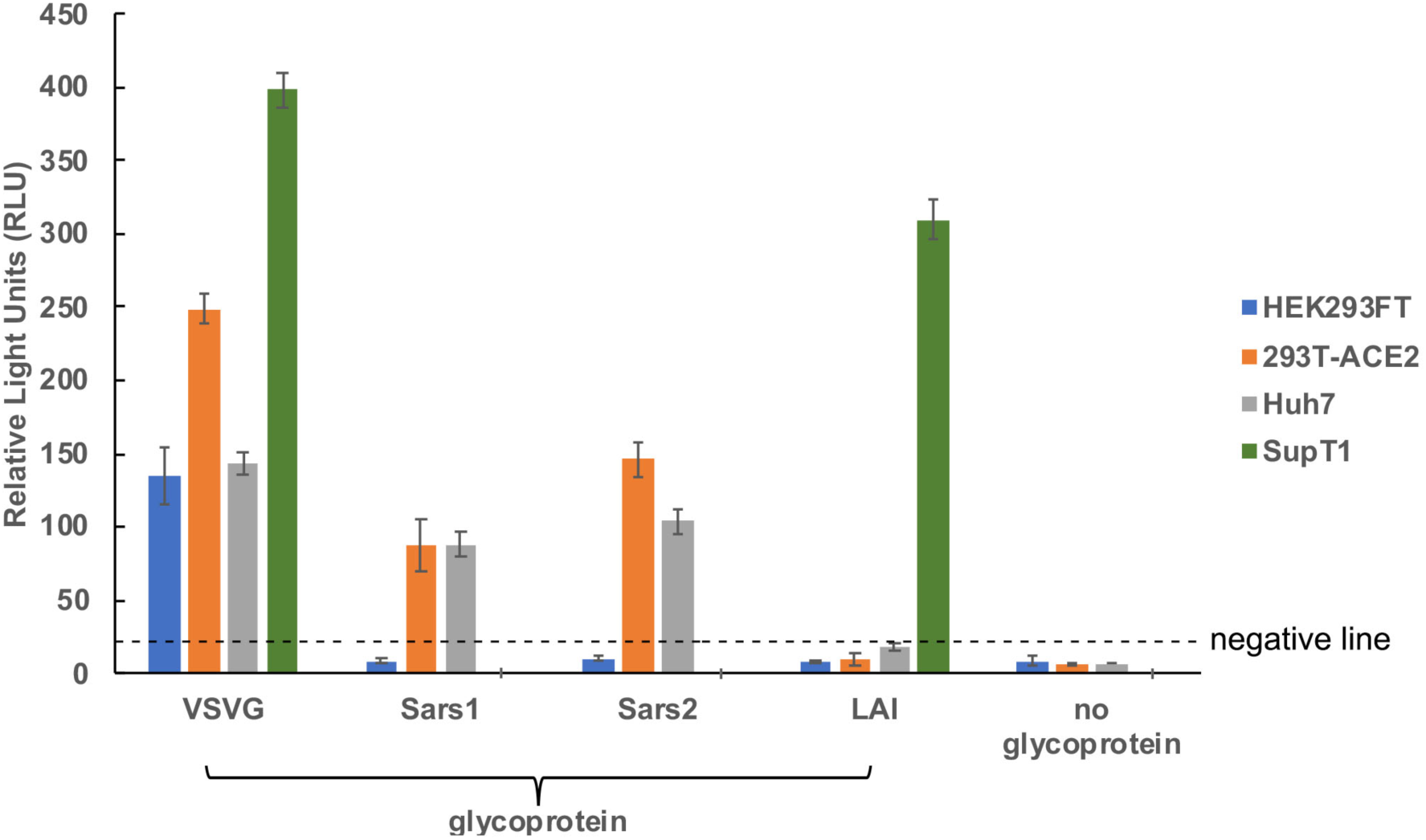
Infectivity of pseudotyped MLV Viruses. SARS-CoV-2 spike pseudotyped MLV viruses as well as VSV-G, SARS-CoV-1 spike, and HIV-1 LAIgp160 pseudotyped MLV viruses were tested on HEK293FT, 293T-ACE2, Huh7 and SupT1 cells. 100μL of undiluted virus (except for VSV-G pseudotype, which was diluted 1:100) was mixed with 100μL of medium and added to cells. Luciferase was measured at 2 days post-infection and values are per well of a 96-well plate. Negative line indicates mean+3SD of luciferase values obtained with virions devoid of glycoprotein.

Since 293T-ACE2 cells showed the highest susceptibility to both SARS-CoV-1 and SARS-CoV-2 pseudotyped MLV viruses, further experiments were all performed in 293T-ACE2 cells. The ultimate goal of our studies was to develop an antibody virus neutralization test based on the above pseudotyped virus. To test for neutralization activity, serum samples from 12 de-identified COVID-19 patients were tested for their ability to neutralize pseudotyped MLV viruses.. Samples 1-6 were plasma obtained from patients who had a SARS-CoV-2 positive test for nucleocapsid-specific IgG (Abbot; samples 1 and 12), spike-specific IgG (Euroimmun; samples 2, 4, 5, and 6) or Nucleic Acid Amplification test (ARUP Laboratories; samples 3, and 8-11) either SARS-CoV-2 nucleocapsid ELISA or SARS-CoV-2 PCR.

As shown in Figure 2, 11 out of 12 patient serum samples showed neutralizing activity against SARS-CoV-2-spike pseudotyped MLV viruses, with neutralizing titers-50 (NT_50_) that ranged from 1:25 to 1:1,417. Eight out of the 12 samples displayed detectable NT_80_. We also tested five historical samples from patients who were hospitalized for severe influenza infection in 2016, all of which tested negative in the neutralization assay (NT_50_ < 25; Figure 3).

**Figure 2.**
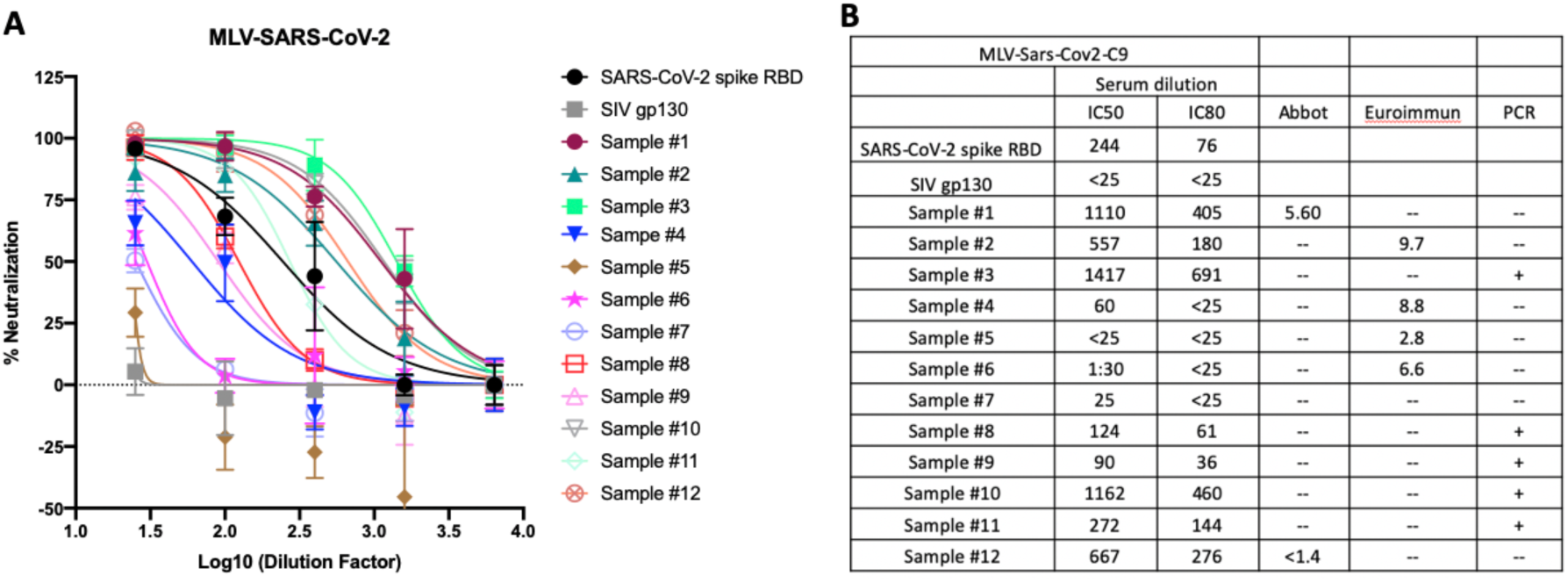
Neutralizing activity of COVID-19 patient serum against and SARS-CoV-2 pseudotyped MLV. A. Serum of COVID-19 patients were pre-incubated with SARS-CoV-2 spike pseudotyped MLV at 37 °C for 1h. Serum and virus mixture were then incubated with 293T-ACE2 cells for 2 days. SARS-CoV-2 spike RBD was used as a positive control. SIV gp130 was used as a negative control. Luciferase was measured to assess infection. Percentage of neutralization was calculated. **B**. Neutralization titer 50 and 80 (NT_50_, NT_80_) were calculated as the reciprocal of the dilution resulting in 50 and 80% neutralization, respectively. ELISA tests by Abbot and Euroimmun are not quantitative.

**Figure 3.**
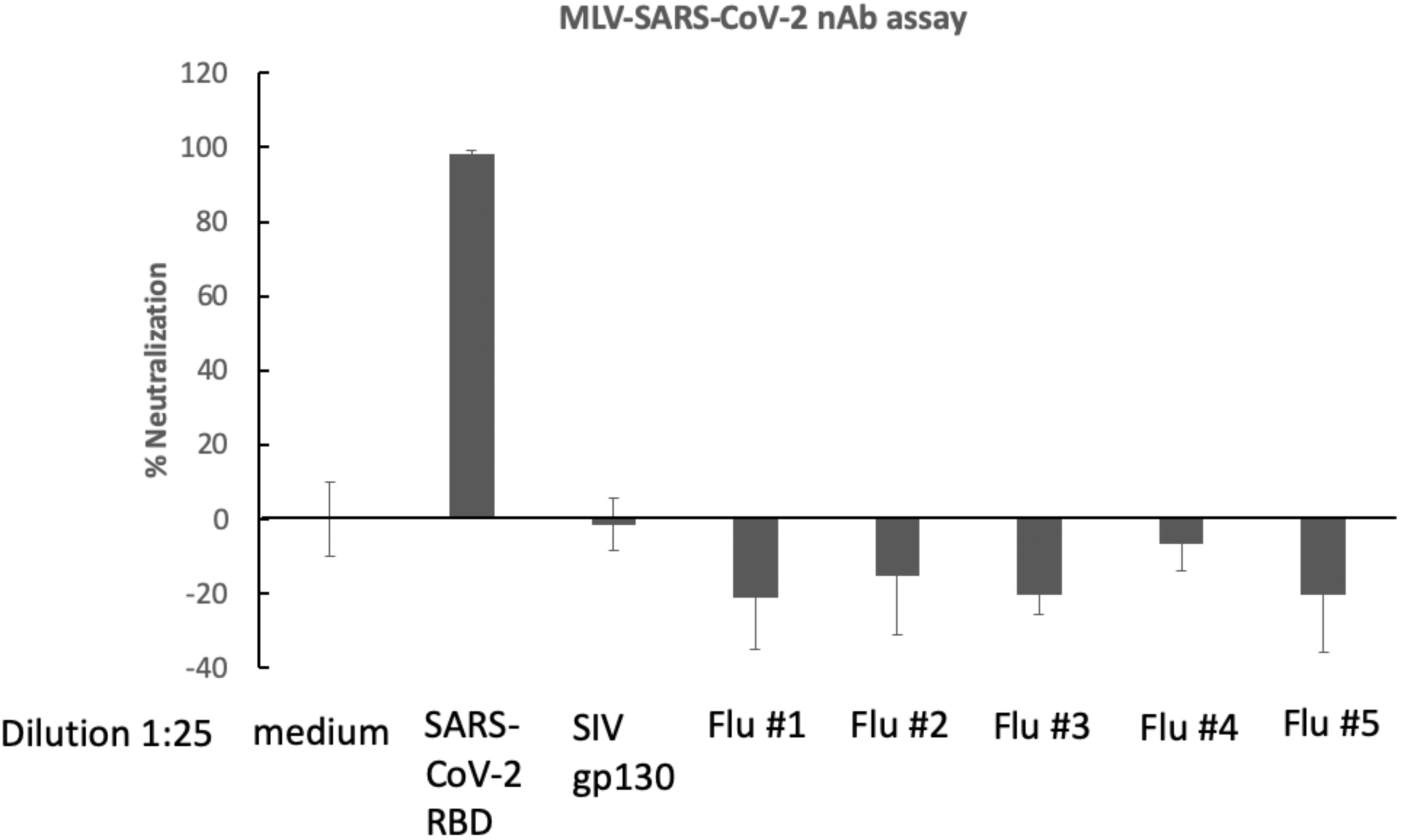
Sera from hospitalized flu patients had no neutralizing activity against SARS-CoV-1 pseudovirions. Cryopreserved serum samples from hospitalized flu patients from 2016 were pre-incubated with SARS-CoV-2 spike pseudotyped MLV at 37°C for 1h. Serum and virus mixture were then incubated with 293T-ACE2 cells for 2 days. SARS-CoV-2 spike RBD was used as a positive control. SIV gp130 was used as a negative control.

To test for specificity of neutralization, we asked whether neutralizing antibodies from SARS-CoV-2 patients would exhibit cross-reactivity against a pseudotype expressing SARS-CoV-1 (Figure 4). We tested samples #1, 2 and 3, which had the highest NT_50_ and NT_80_. None of these sera had detectable neutralizing activity (NT_50_ <25) against the SARS-CoV-1 pseudotype, which is consistent with previous reports [11-13].

**Figure 4.**
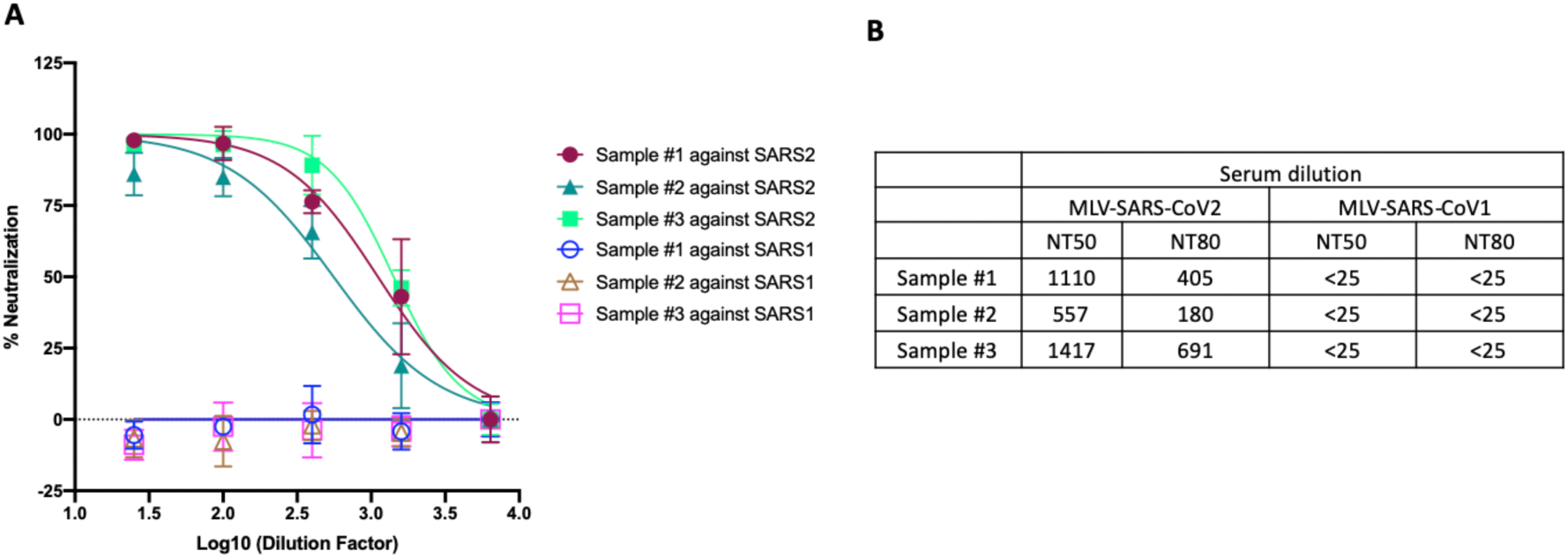
Neutralizing activity of COVID-19 patient serum against SARS-CoV-1 pseudovirions. (**A**) Serum from COVID-19 patients were pre-incubated with SARS-CoV-1 spike pseudotyped MLV at 37 °C for 1h. Serum and virus mixture were then incubated with 293T-ACE2 cells. Percentage of neutralization was calculated. (**B**) NT_50_ and NT_80_ were calculated as above. Samples #1, 2 and 3# were tested previously (Figure 2).

As a positive control and also as a standard to monitor variability between neutralization experiments, we used recombinant soluble receptor binding domain from SARS-CoV-2 spike protein. We produced this protein via transient transfection in HEK293FT cells using a mammalian expression vector (pCAGGS) encoding amino acids 319 to 542 of from SARS-CoV-2 S1, encompassing the RBD (BEI Resources, cat.# NR-52309). The apparent NT_50_ of RBD against SARS-CoV-2 / MLV pseudotype was 1:244. As a negative control for neutralization, the surface glycoprotein from the simian immunodeficiency virus, SIVmac gp130 [10], was similarly produced by transfection.

### Conclusions

In summary, we have developed a simple and rapid assay based on pseudovirion particles, which should allow for specific measurement of neutralizing titers in plasma against SARS-CoV-2 in the context of biocontainment level 2 laboratories. Using SARS-CoV-2 RBD as a control in this assay, we observe that, as expected, SARS-CoV-2 RBD was able to block infection with SARS-CoV-2.

## DECLARATIONS

### Ethics approval and consent to participate

We used de-identified, archived plasma or serum samples throughout the study.

### Consent for publication

All authors have agreed to publication of the manuscript.

### Availability of data and materials

Plasmid constructs and methodology are available upon request. Aliquots of plasma and serum are in limiting quantities and may be available depending on amount requested.

### Authors contributions

YZ, ETL, EAI, ESCPW conducted experiments. JL, IC, MC, AMS and VP designed the study. JCD, PS, JR and MTR selected and contributed archived samples.

## Acknowledgments

The construct pcDNA3.1-SARS-2-S-C9 was a generous gift from Tom Gallagher at Loyola University. Human Codon Optimized HEK293T-hACE2 cells were kindly provided by Adam Bailey and Emma Winkler in the laboratory of Michael Diamond at Washington University in St. Louis School of Medicine. We also wish to thank Eloisa Yuste for helpful technical suggestions.

## Abbreviations

SARS: severe acute respiratory syndrome
SARS-CoV-2: severe acute respiratory syndrome coronavirus 2
COVID-19: coronavirus disease 2019
DMEM: Dulbecco’s modified Eagle’s medium
FBS: fetal bovine serum
RBD: receptor binding domain
NT: neutralizing titer
MLV: murine leukemia virus
VSV-G: vesicular stomatitis virus glycoprotein
HIV: human immunodeficiency virus
ACE2: Angiotensin Converting Enzyme 2

